# The VVBlue assay: a plate-readable, dye exclusion-based cell viability assay for the toxicological testing of chemicals

**DOI:** 10.1101/2024.08.27.609841

**Authors:** Marianne Vitipon, Esther Akingbagbohun, Thierry Rabilloud

## Abstract

A viability test for in vitro cultures, based on the intake of the textile dye Alphazurine A by dead cells and its exclusion by viable cells, is described. This test uses the affinity of Alphazurine A for proteins, so that the dye is retained in dead cells even after rinsing, while its anionic character prevents it from entering live cells. This feature makes this dye exclusion test amenable to a reading in a plate format. The Alphazurine viability test provides an indicator of the absolute number of dead cells present in the culture well. To reach a cell viability index, a “dead cells” control (e.g. cells killed with ethanol) must be added. We also describe a double viability test, which first uses the Alphazurine assay to provide the number of dead cells then a crystal violet assay to provide an index of the number of cells present in the plate. This double test provides a complete appraisal of the situation in the cell culture wells, and has been compared to other viability tests such as propidium iodide exclusion or tetrazolium reduction. Its performances to study the toxicity of substances such as pigments are also established, and allowed us to publish the first public toxicological data on the recently described Pigment Blue 86.

## Introduction

There is more and more pressure on the chemical industry to market safe products for the public consumers, and products with documented toxicology for professional customers. In some part of the world, e.g. European Union, it has become mandatory to document the toxicology of every product marketed over 1 ton/year. However, this assessment relies on regulatory tests that are both long and expensive. This poses a problem to the industry, as the whole development of a new chemical may be canceled at the very end of the process by the toxicology assessment. There is thus a need of simple, fast and cheap toxicological tests to perform early toxicological tests during the development cycle of new products, to comply to the “fail early, fail cheap” principle that has long been adopted in the pharmaceutical industry.

To this purpose, in vitro toxicological tests combine speed and relatively low costs, and are often amenable to automation, which allows many tests to be performed and to test several conditions simultaneously.

Within this frame on in vitro toxicology, the determination of cell viability is an operation performed routinely in all cell biology laboratories. While performed at a relatively low scale for routine cell culture maintenance, it becomes crucial in certain areas of biology such as toxicology, where the determination of cell viability is an essential foundation for all subsequent analyses.

Depending on the cellular parameter used to determine it, cell viability assays can be classified as direct or indirect assays. Direct assays rely on the contrast between the membrane permeability existing for dead cells compared to the much more tightly-regulated permeability observed on live cells. Thus, direct cell viability assays use either the permeability of dead cells to certain dyes or fluorescent probes added in the medium or the release of intracellular components in the medium. Historically, the contrast between the impermeability of live cells to anionic dyes (and multicharged cationic dyes) and the permeability of dead cells to these dyes has been used to build cell viability assays, the forst dye used to this purpose being Trypan blue [1]. When used in a classical microscopy format, dyes in addition to Trypan blue, such as Eosin, Erythrosin and Safranin have been used and compared [2]. However, these dyes have various properties, e.g. in their toxicity or speed of penetration into dead cells that limit their application for large scale experiments. Furthermore, they rely on human interpretation and are thus operator dependent, although automated solutions now exist when working with classical dyes such as Trypan blue. In a flow cytometry format, which is better suited for medium-large scale experiments and is less prone to operator dependency, propidium iodide is used as a fluorescent probe to discriminate between live and dead cells [3]. However, this format is still a serial operation, requiring an expensive equipement and is prone to some artifacts, e.g. strongly negatively charged particles that bind propidium iodide can be misidentified as cells during gate analysis. Moreover, the definition of the gates to discriminate between live and dead cells is not necessarily obvious, as a fluorescence continuum is often observed between the two main clouds of points.

In addition, for these optical methods requiring cell observation or measurement of cellular fluorescence, the cells must ideally be transparent. Cells that are rendered opaque, e.g. by internalization of dark nanoparticles, cannot be tested by such methods.

In the other direction, i.e. the release of intracellular components into the extracellular medium, the only test that has gained popularity is based on the release of lactate dehydrogenase, i.e. a very abundant cytosolic enzyme that is relatively easy to assay [4]. However, this assay still relies on the quantitative determination of an enzyme activity, i.e. a process that requires careful time and temperature control, and that can be biased if the extracellular medium contains substances interfering with the assay, which is sometimes encountered in toxicology experiments. This phenomenon has been described for metal ions, for example released by nanoparticles [5,6], but also for bacteria [7], and for nanoparticles that adsorb LDH [8].

These limitations have stimulated the development of indirect assays that are substantially easier to automate, and thus less prone to inter-operator variations. Depending on the cellular parameter tested, several classes of indirect viability assays can be described.

The first and most popular is based on the cellular redox metabolism and the existence of the pool of reduced dinucleotides (NADH and NADPH). Due to their low redox potential, these reduced dinucleotides can convert either weakly colored tetrazolium salts into strongly colored formazans [9,10] or convert non fluorescent molecules into fluorescent ones [11]. Although extremely popular, these assays are not devoided of caveats. The first one is linked to their indirect nature. For example, a 50% reduction in formazan production in treated cells compared to controls does not tell whether half of the cells are dead and half of them have a completely normal metabolism, or whether all cells are alive but with a two-fold reduced metabolism. As an example, bromopyruvate, which is a metabolic poison, gives erratic results in tetrazolium and resazurin reduction assays [12]. Other caveats encountered with such assays in toxicology experiments can be due to the intrinsic redox capacity of the substances to be tested [13], or to the superoxide production by the cells in response to the experimental conditions [14]. This may lead to “viabilities” over 100% when such an interference or cell activation is encountered [13,14], which of course cannot happen. Further problems are encountered when the substance to be tested is a highly hydrophobic particle able to strongly adsorb the reduced molecule used as an indicator of cell metabolism, either formazans [15] or resorufin [16]. Furthermore, the resazurin reduction assay is prone to a specific artifact represented by the over-reduction of the desired fluorescent product resorufin, into the non-fluorescent product dihydroresorufin [11].

A second indirect cell viability assay relies on the proton pumping activity of lysosomes. As this phenomenon is ATP-dependent, it quickly stops after cell death. This phenomenon leads in turn to the ability of certain cationic dyes, such as neutral red, to be accumulated into lysosomes, in live cells only. This has led to the development of the neutral red uptake assay as an indirect cell viability assay [17]. This assay is however also prone to biases, arising for example from light-absorbing compounds [18], from selective inhibition of lysosomal proton pumping in viable cells [19], or from cell activation. In the latter case, “viabilities” over 100% can be observed [20].

Other indirect cell viability assay relies on cell number, which is not determined in an absolute way but rather via cellular macromolecules. Such assays include the crystal violet assay [21], which stains nuclei, and the sulforhodamine assay [22], which estimates the protein content. The rationale under these assays is that dead cells will not attach and/or detach from cell culture supports. In turn, these assays will apply only to adherent cells, which however represent the vast majority of cultured cells. Quite unfortunately, the underlying hypothesis of these assays on dead cells detachment has been shown to be incorrect in some cases [23].

Due to these various limitations, rather than relying on indirect assay multiplexing [24], we sought to develop an assay that would rely on dye exclusion, but would not need either microscopy or flow cytometry, and would rather be readable in plate format. This would require a staining sequence similar to the one used for crystal violet or neutral red staining, i.e. staining with the dye, rinsing to eliminate the excess dye that would prevent reading in the plate reader, then dye elution to read the absorbance from the culture supports. When classical exclusion dyes or even sulforhodamine B [2,22] were tested in such an approach, no signal was detectable after rinsing, probably due to the insufficient affinity of the dyes for intracellular components, thereby not withstanding the rinsing phase. Further substantiation of this hypothesis was obtained from a variation of the dye exclusion protocol using the fluorescence of erythrosin B in dead cells [25]. This protocol does not use any rinsing step, but rather uses a very low probe concentration (5µM) to limit the background fluorescence, thereby allowing sufficient contrast to identify the dead cells by photographic fluorescence light accumulation. As this method appeared as not suitable for medium-large scale experiments because of the need for a photographic process, we decided to test several dyes in order to find one that would not penetrate live cells, and have sufficient affinity for cellular components to allow for a stain-rinse-elution sequence.

In addition, this process should allow to develop a test that should be insensitive to optical interferences, and thus suitable for the toxicological testing of even the most difficult chemicals in in vitro toxicology, such as pigments, which are both particular in nature of optically opaque.

## Material and methods

### 2.1. Materials

All dyes were purchased from Sigma at the highest purity available. Zinc oxide (<100 nm, catalog number # 721085) and silver (<100nm, catalog number # 758329) nanoparticles dispersions were purchased from Sigma. Ethanol (96%) was purchased from RSE, acetic acid (analytical grade) from Sigma. Pigment black 33 (mixed manganese iron oxide #47501) and Pigment Bue 86 (Yttrium manganese indium oxide #453208) were purchased from Kremer pigments, and Pigment Black 14 (manganese dioxide) was purchased from Zecchi (reference C0999). The pigments were suspended in gum arabic (100 mg/ml, used as an anti-agglomeration reagent) that had been previously sterilized overnight at 80°C. Then, the suspensions of pigments (100 mg/mL) was sterilized overnight again before being was sonicated in a Vibra Cell VC 750 sonicator equipped with a Cup Horn probe (VWR, Fontenay-sous-Bois, France) with the following settings: time = 30min – 1 second ON, 1 second OFF – Amplitude = 60%, corresponding to 90 W/pulse.

Before adding to the cell culture, nanoparticles dispersions were diluted in sterile deionized water.

### 2.2 Cell culture

The mouse macrophage cell line J774A.1 and the alveolar epithelial cell line A549 were obtained from the European Cell Culture Collection (Salisbury, UK). The cells were cultured in DMEM medium + 10% fetal bovine serum (FBS). For routine culture, J774 cells were seeded on non-adherent flasks (e.g. suspension culture flasks from Greiner) at 200,000 cells/ml and harvested 48 hours later, at 1,000,000 cells/ml. A549 cells were seeded on cell culture adherent flasks at a cell density of 10,000 cells/cm^2^ and split by trypsinization every 3 days.

For treatment with particles, the following scheme was used: J774 cells were first seeded at 400,000 cells/mL into 24-well adherent plates (FALCON®, Corning Incorporated) or 12-well non-adherent plates (CELLSTAR®, Greiner), depending on the viability test performed. They were exposed to zinc oxide or silver nanoparticles or silver nitrate or pigments on the following day and harvested after a further 24h in culture. In some experiments, the cells were cultured in DMEM medium + 1% fetal bovine serum (FBS).

For A549 cells, they were first seeded at 100,000 cells/ cm^2^ into 24-well adherent plates (FALCON®, Corning Incorporated). They were exposed to silver nitrate on the following day and analyzed after a further 24h in culture.

Cells were used at passage numbers from 5 to 15 post-reception from the repository.

### 2.3. Viability assays

#### 2.3.1. Classical assay: propidium iodide exclusion assay

Propidium Iodide cell viability was measured using FACScalibur flow cytometer equipped with CellQuest software (6.0, Becton Dickinson Biosciences, Le Pont-de-Claix, France). After detachment from the culture plastic support by repeated flushing, cells were dyed with propidium iodide (480 nm excitation and 600 nm emission) at 1 μg/ml final concentration for 5 minutes prior to analysis.

#### 2.3.2. Classical assays: WST1 and MTT metabolic assay

The metabolic activity of cells was evaluated by using the cell proliferation reagent WST-1 (Sigmaaldrich, #11644807001). Briefly, after 24 hours exposure to chemicals, plates were centrifuged for 3 min at 1200 rpm. Subsequently, the culture medium was removed from the test conditions, while for the “dead cells” control condition the cells were treated with ethanol (50% v/v) for 3 min. After rinsing all the wells with a DMEM without phenol red (PAN Biotech, #1190124), WST-1 reagent was added at a final concentration (1:50) in phenol red-free DMEM and left incubate at 37°C 5%CO2 for 20 min and 10-12 min, for J774A.1 and A549 respectively. Absorbances were measured at 440 nm with a plate reader (Infinte M200, Tecan, Durham, NC, USA).

In other experiments, metabolic activity of cells was evaluated by using the MTT assay (Sigmaaldrich, #M5655). Briefly, after 24 hours exposure to chemicals, the culture medium was removed from the test conditions. The medium was replaced by DMEM without phenol red (PAN Biotech, #1190124), to which 10 µl of a 20mg/ml MTT solution in absolute ethanol had been added. The cells were returned to the culture incubator for 1 hour. The culture medium was then removed and the precipitated formazan was eluted with a solution of 90% ethanol, 9% water and 1% acetic acid (all by volumes) for 45 min with agitation. Absorbances were measured at 570 nm with a plate reader.

#### 2.3.3. the VVBlue assay

The VVBlue assay was conducted according to the following protocol detailed in **Supplementary Document 1**. Unless otherwise indicated, the volume used for all protocol steps is equal to the volume of culture medium during cell culture. Prior to staining, cells were centrifuged 3 min at 1200 rpm to collect both adherent and detached cells. Subsequently, the culture medium was replaced with fresh medium for the tested conditions, while cells in the “dead cells” control conditions were treated with ethanol (50%v/v) for 3 to 5 min. After washing the “dead cells” control conditions with fresh medium, alphazurine A was added (from a stock solution at 10 mg/ml in water) directly to the cell culture medium at a final concentration of 0.2 mg/ml, followed by a 40 min incubation at 37°C. Three rinses in PBS were performed to remove excess dye, followed each time by a 3 min centrifugation at 1200 rpm. The cell-associated dye was then eluted with a solution of 50% ethanol 1% acetic acid for 30 min with agitation. A volume equal to the one of initial culture medium was used. After centrifugation, a fraction was transferred into a new plate, and absorbance was measured at 637 nm. The remaining eluate was removed, and cells were rinsed with PBS to rehydrate them before adding a volume of crystal violet solution at 4 µg/mL and incubated for 30 min at room temperature. Three rinses in PBS were performed to remove excess dye, followed each time by a 3 min centrifugation at 1200 rpm. The cell-associated dye was then eluted with a solution of 50% ethanol 1% acetic acid for 45 min with agitation. After centrifugation, a fraction was transferred into a new plate, and absorbance was measured at 590 nm.

When optical interference was expected, as in the case of pigments, a second series of controls was implemented. These controls consisted of plates devoid of any cells, but treated with the same concentrations of pigments used in the toxicological testing. These plates were then submitted to the VVBlue test and the obtained absorbances deduced from the ones obtained in the cells-containing plates.

### 2.4. Microscopy visualization experiments

After the adequate treatments, the culture medium was replaced by phenol red-free DMEM for imaging. Images were taken before and after the alphazurine treatment, including the PBS washes, but before any elution step. The cells were visualized under an optical microscope (Motic AE21) set at x10 magnification. The images were captured with an Axiocam 105 color camera and processed uniformly using ImageJ to standardize contrast and saturation levels.

## Results

### 3.1. Dye selection

In order to select suitable dyes, we performed preliminary experiments in a simplified format. Cells were grown in 6 well plates, and in one half of the wells the cells were killed with a lethal dose of silver nanoparticles. All wells were then treated with 0.2 mg/ml dye dissolved in culture medium for 30 minutes. The dye solution was then removed, the wells rinsed twice with PBS, and the dye was eluted with an acidic ethanol solution (1% acetic acid, 50% ethanol). The absorbance of the wells corresponding to live and dead cells was then compared. The following dyes were tested through this method: erythrosin (C.I. 45430), crystal violet (C.I. 42555), methyl green (C.I. 42585), brilliant green (C.I. 42040), safranin (C.I. 50240), methylene green (C.I.52020), ponceau S (C.I. 27195), light green SF (C.I. 42095), acid violet 17 (C.I. 42650), erioglaucine (C.I. 42090), alphazurine A (C.I. 42080), Coomassie blue G250 (C.I. 42655), copper phthalocyanine tetrasulfonate, lissamine green (C.I. 44090), thioflavin S (C.I. 49010), thiazole yellow (C.I. 19540), Nile blue (C.I. 51180) and naphthol yellow (C.I. 10316).

Besides classical exclusion dyes such as erythrosin and safranin, other dyes were selected on the basis of their use as wool (protein) dyes (light green SF, acid violet 17, erioglaucine, alphazurine A, Coomassie blue G250 thioflavin S, thiazole yellow and lissamine green), as nucleic acid dyes (crystal violet, methyl green, brilliant green) or as protein stains in blotting (Ponceau S, copper phthalocyanine tetrasulfonate).

Of all those, only the textile dyes acid violet 17 and alphazurine A offered an interesting absorbance difference between live cells and dead cells, as shown in **Fig 1**. Coomassie blue did stain dead cells well but appeared too hydrophobic, as its excess was not easily removed from the culture wells. Acid violet 17 also gave a rather high absorbance in the wells with live cells. Hence, we finally focused on the alphazurine A dye.

**Fig 1.**
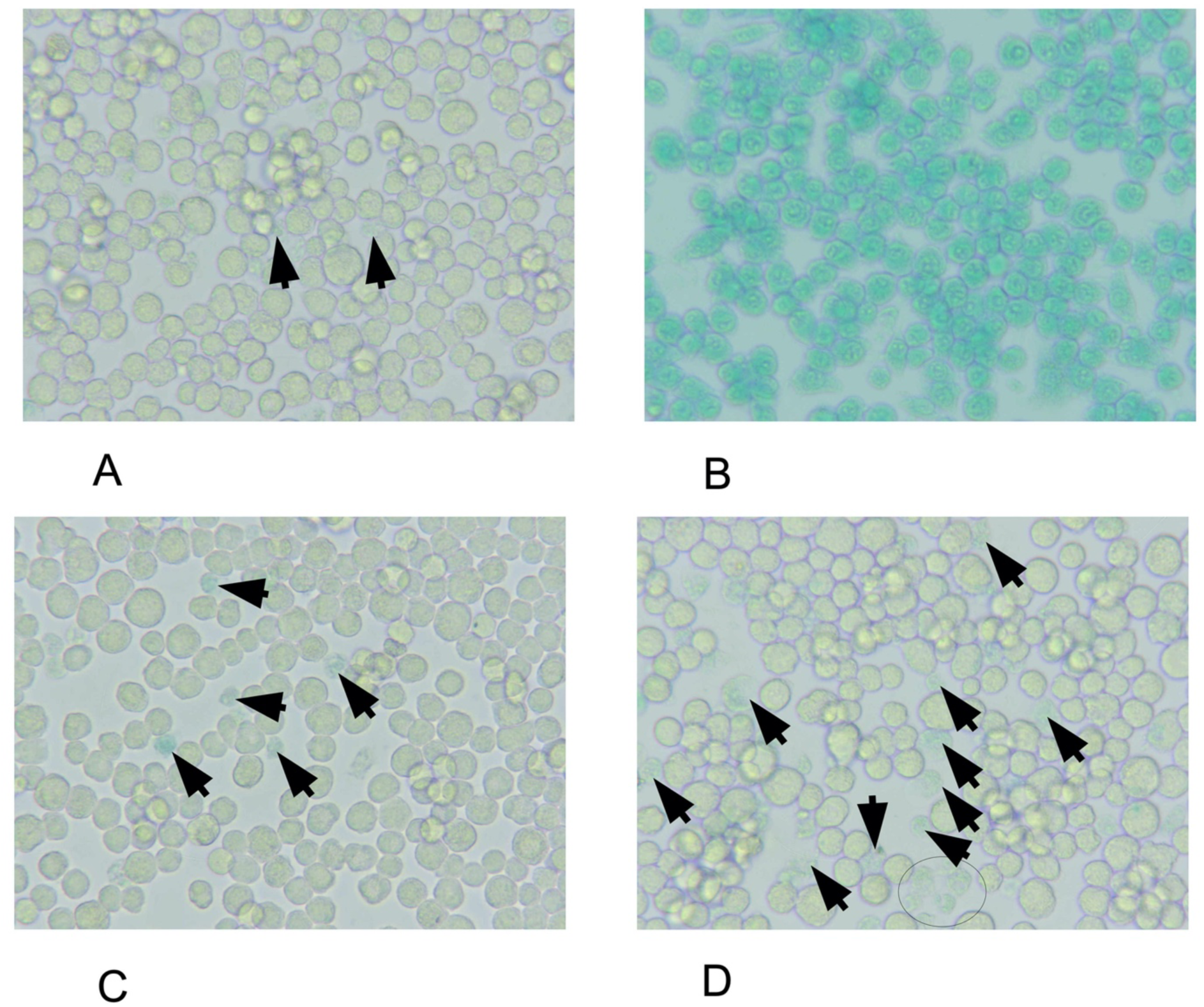
Microscopy viewing of live and dead cells after treatment with alphazurine. **A** J774A.1 cells were seeded at a density of 500,000 cells/mL in 24-well adherent plates containing DMEM supplemented with 1% horse serum and 1% streptomycin-penicillin. Cells were either kept untreated (panel A), killed with 50% ethanol for 5 minutes (panel B) or treated with 10 or 20µg/ml silver nitrate for 24 hours (panels C and D). The black arrows and ellipses in the panels show the dead cells occurring in the various conditions

### 3.2. Assay development

We first optimized the rinsing and elution phases. For rinsing, a buffer saline solution (i.e.with an ionic strength equivalent to 150mM sodium chloride) was tested. Alternatively, a low ionic strength solution containing 10mM Hepes pH 7.5, 1mM MgCl2 and 300mM sucrose was also tested. The rinsing efficiency was visually evaluated by the retention of the green color in the wells containing 100% dead cells wells, while the green color should have been removed in the wells containing live cells. Since both solutions performed equally well, the saline solution was selected for its better shelf life.

For elution, we tested either an acidic solution (1%acetic acid, 50% ethanol) or a basic solution (20mM Tris base in 50% ethanol), as basic solutions have been used to elute triphenylmethane dyes from stained polyacrylamide gels bands [26,27]. However, basic elution proved inappropriate, as evaluation by microscopy showed that it removed the cell layer from the plates, which prevented the construction of a robust assay.

We then investigated the linearity of the assay. For this purpose, a defined number of cells were seeded in 24-well plates, let to adhere for 1 hour, and then killed by the addition of 50% ethanol. After rehydration of the dead cell layers in DMEM, the cells were stained with alphazurine, rinsed and the dye eluted with the acid ethanol solution. The absorbance at 637 nm was recorded, and the results are shown in **Fig 2A**. The results show that this calibration curve is linear in the 0-05 OD range, corresponding to cell numbers up to 1 million cells per well in a 24-wells plate, and that the method indeed indicated the number of dead cells present in the wells. However, it could not be excluded that in some experiments dead cells may detach, or that depending on the experimental conditions, the adherence of cells may vary. To normalize the number of dead cells (alphazurine-positive) by the total number of cells present in each well, we used an assay that is based on cell numbers as a second step, i.e. the crystal violet assay, as described in [28]. When applied to the wells with defined cells numbers after the alphazurine staining and elution, the results, described in **Fig 2B**, showed that the crystal violet signal correlated well with the cell numbers seeded in the plates, with a correlation coefficient of 0.96.

**Fig 2.**
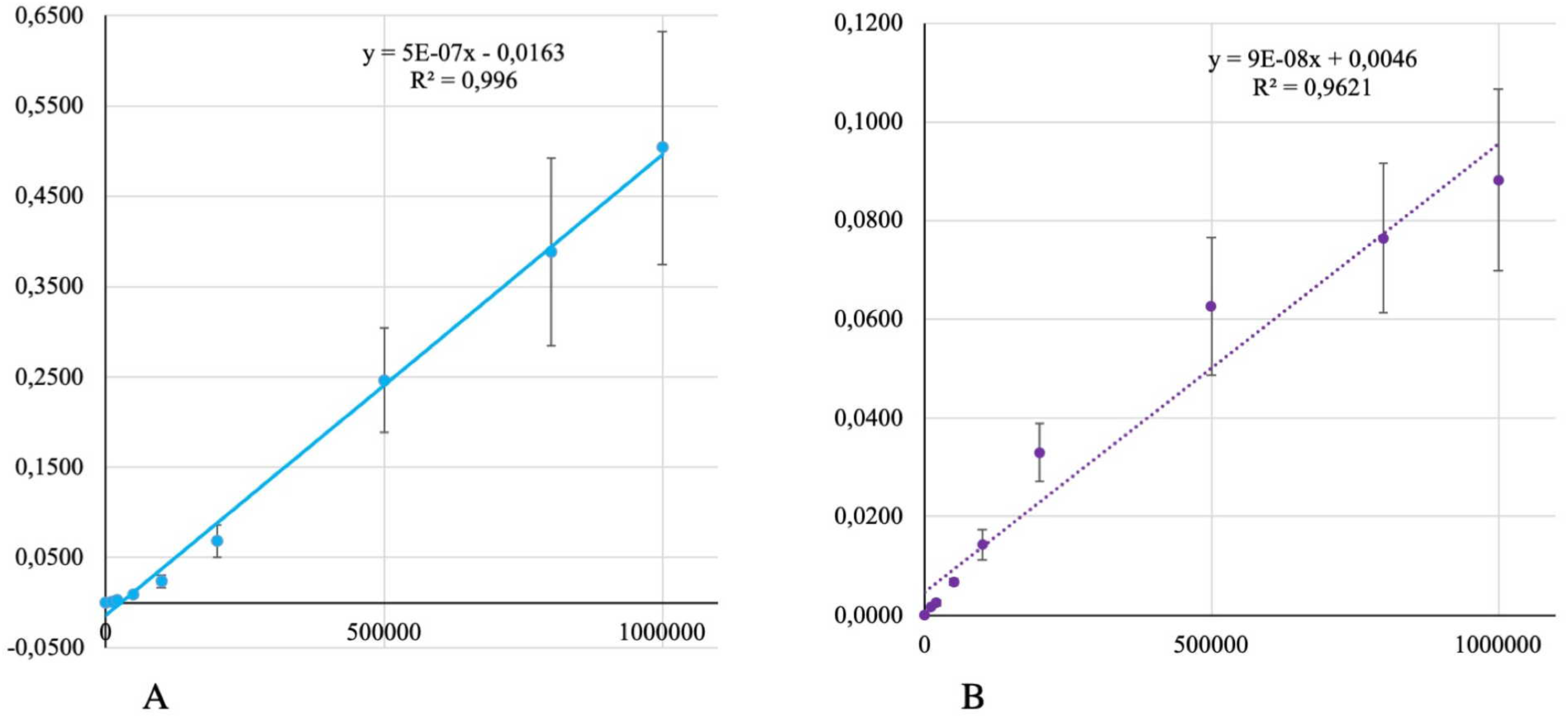
Relationship between J774A.1 cell numbers inoculated into 24-well plates and absorbance after staining. (A) Staining with alphazurine A in DMEM (B) Staining with crystal violet. The absorbance was measured at 637 nm for alphazurine A and 590 nm for crystal violet. Bars indicate SD of readings from triplicate experiments.

When several experiments were performed, a variability in the absorbance values was observed, as shown in **Fig 3**.

**Fig 3.**
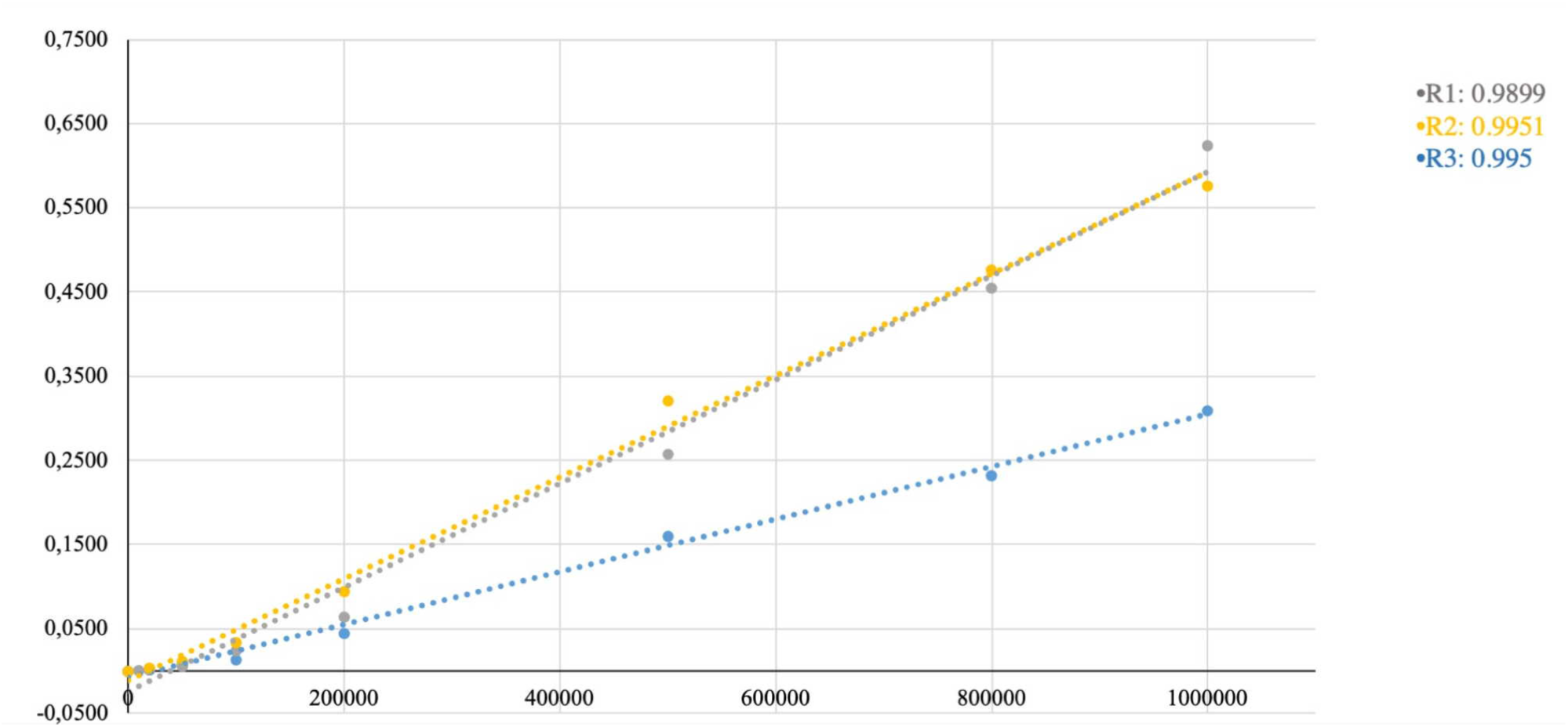
Inter-experiment variability of the relationship between J774A.1 cell numbers inoculated into 24-well plates and absorbance after staining with alphazurine A in DMEM. Three independent experiments were conducted. In each experiment, a linear correlation was observed, with a R² coefficient of 0.9899 for the grey dots, 0.9951 for the yellow dots, and 0.995 for the blue dots.

Consequently, the utilization of a single numerical equation that would convert the OD signals directly into cell numbers is impossible. However, within the confines of a single experiment, linearity is upheld with a high R² coefficient as shown in Figure 3. Thus, the incorporation of additional control points in each experiment is crucial to use the linearity correlation observed for these two dyes. .

For the Alphazurine stain, a “cells only” control is imperative to ascertain the initial viability of the cells and to relate to the theoretical cell count in the wells. Additionally, a “cells only 100% dead” control is essential. These two controls enable the establishment, within the same experimentation, of the 0% dead cells and 100% dead cells points found in the standard ranges, facilitating the subsequent application of linearity correlation. Another control condition needed is the “empty well”, which have undergone the same coloration steps as the others conditions as outlined in Table 1, to be used as a blank.

Once all the absorbance have been obtained, they are processed as follows to deduce cell viability. The absorbances resulting from these stains were normalized against the blank to eliminate plastic and solvent absorbance. Subsequently, for crystal violet, a normalized absorbance ratio was calculated between the absorbance of each condition and the average absorbance of the control conditions, where the cell count remains unchanged, serving as the reference. This index gave the proportion of cells still present in the plate at the end of the assay.

The alphazurine data were then used. The background-corrected absorbance were used to calculate the absorbance due to dead cells in every well. The average absorbance of the ethanol-killed cells and the average absorbance of the control cells were used to determine the absorbance linked to 0% viability and 100% viability, respectively, leading to the calculation of the absorbance linked to dead cells in each well. The substraction of this value to the crystal violet value led to the proportion of live cells. Theses calculations are detailed in **Supplementary Document 1**.

### 3.3. Cell viability assessments comparisons: VVBlue vs. Propidium Iodide and WST1 Assays

To ensure the robustness of our assay, we conducted a comparative analysis of J774A.1 cell viability using the VVBlue assay against a well-established cell viability assay employing propidium iodide (PI). Cells were seeded at a density of 400.000 cells/mL in a 24-well adherent plate for VVBlue assay, and, in a 12-well non-adherent plate for PI assay. Cells were exposed to increasing concentrations of particles, either zinc oxide (ZnO) or silver nanoparticles (AgNP), on the following day and harvested after 24h of exposure to particles.

For the PI cell viability assay, cells were pre-stained with propidium iodide at 1 μg/ml in PBS for 5min at RT, washed with PBS before analysis using flow cytometry. The cell viability under different conditions was normalized to the control group composed of cells without particles. As for the VVBlue assay, the samples were processed as explained previously in Table 1. The obtained results are illustrated in **Fig 4**.

**Fig 4:**
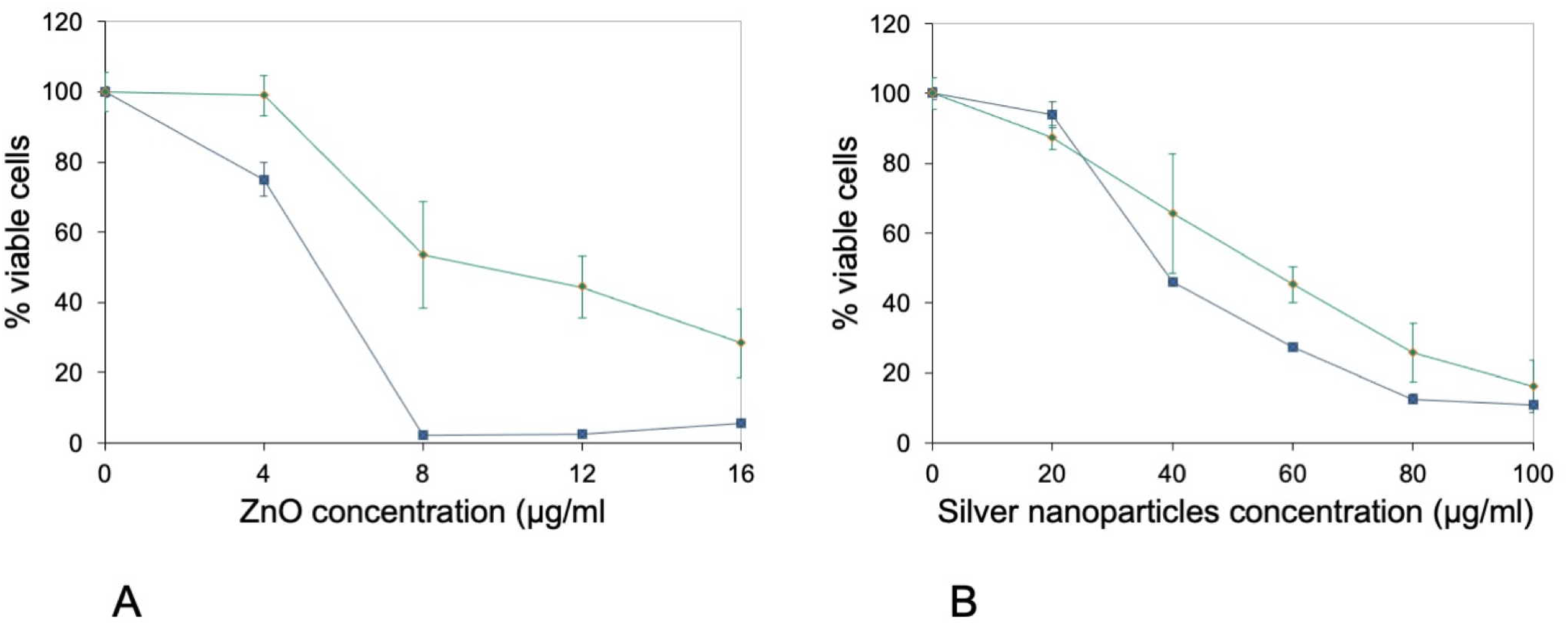
Comparison of the propidium iodide and VVBlue assays. A) Cell viability after 24 hours of exposure to zinc oxide nanoparticles Blue curve: viability measured by the propidium iodide assay (cells cultured on non-adherent plates). Green curve: viability measured by the VVBlue assay (cells cultured on adherent plates). Experiments were conducted in quadruplicate and represented as mean ± standard deviation B) Cell viability after 24 hours of exposure to silver nanoparticles. Blue curve: viability measured by the propidium iodide assay (cells cultured on non-adherent plates). Green curve: viability measured by the VVBlue assay (cells cultured on adherent plates). Experiments were conducted in triplicate and represented as mean ± standard deviation

Remarkably, the results displayed in **Fig.4** showed similarities between the viability obtained from these two assays. Specifically, upon exposure to zinc oxide, a notable decline in cell viability was evident, particularly above the concentration threshold of 8µg/mL. Similarly, when exposed to silver nanoparticles, a consistent trend in cell viability was observed in both assays. These findings highlight the robustness and reliability of the VVBlue assay, allowing to obtain similar results as the PI assay. However, it should be noted as well that the curves do not superimpose. This may be due to the fact that the cells tested by the VVBlue protocol do not sustain any disturbance in their analysis, while cells tested by the PI protocol must be dislodged from their culture support prior to analysis.

Another popular viability assay is represented by the metabolic assays, e.g. the tetrazolium/formazan assays. We thus performed a comparison of the VVBlue assay with the WST1 reduction assay [29]. To this purpose, we used J774A.1 cells exposed to silver nitrate as the toxicant. Cells were seeded at a density of 400.000 cells/mL in a 24-well adherent plate for both assays. Cells were exposed to increasing concentrations of silver nitrate on the following day and assayed after 24h of exposure to the toxicant. The results, displayed on **Fig 5a**, showed a difference in the LD50 obtained by the two methods. The LD50 determined by the WST1 method is around 37µg/ml, while the LD50 determined by the VVBlue assay was around 160µg/ml. Such a phenomenon has already been described in the literature (e.g. in [30]) and will be discussed later.

**Fig 5:**
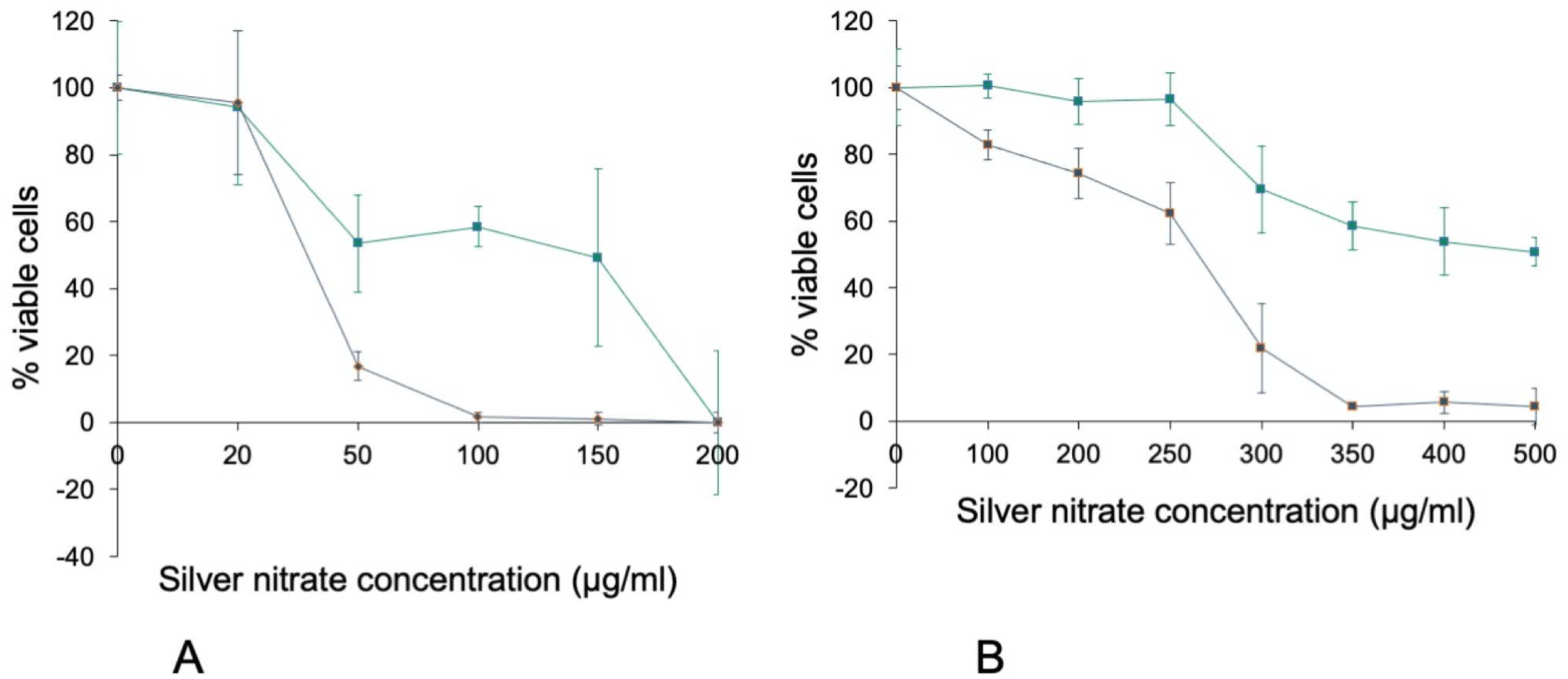
Comparison of the WST1 and VVBlue assays. Cell viability curves after 24 hours of exposure to silver nitrate. Green: VVBlue assay. Blue: WST1 assay. Experiments conducted in quadruplicates and expressed as mean ± standard deviation. Panel A: J774A.1 murine macrophage cell line Panel B: A549 human alveolar epithelial cell line

In order to test our assay on a different cell type, we repeated the WST1 vs. VVBlue assay comparison on a different cell type, i.e. the A549 epithelial cells, still using silver nitrate as a toxicant. The results, displayed on **Fig 5b**, showed once again an important difference in the viabilities determined by the two assays. This figure also showed the interest of the combined alphazurine A/crystal violet assay. While the alphazurine step showed a relatively low number of dead cells, this low number was explained in part by the fact that some of the dead cells detached, a phenomenon taken into account by the crystal violet step. The combination allowed a much more precise determination of the real numbers of viable cells remaining at the end of the assay, and thus of the toxicity.

### 3.4. Performance of the VVBlue under severe optical situations

The VVBlue assay is also performing well under severe situations where other dye-exclusion assays become impracticable. A typical case is the determination of the toxicity of pigment particles, which are opaque, highly colored, and often poorly toxic. These features rendered conventional tests, such as PI or Trypan blue, unsuitable for accurate toxicity assessment. In this precise experimental setup, cells were initially seeded at 500,000 cells/mL in DMEM supplemented with 1% horse serum for 24 hours, and subsequently exposed to pigment black 33 for an additional 24 hours period. The VVBlue assay was employed for viability determination, facilitating the discrimination between live and dead cells, even exposed to elevated concentration of particles, as shown in **Fig 6**. Notably, unexposed live cells (live cells only) exhibited a distinct refractivity and appeared uncolored (Fig. 6A), whereas unexposed dead cells lost their refringence and retained a blue alphazurine staining (Fig. 6B). Interestingly, the numerous black particles did not impede the accurate distinction of dead cells within the exposed cells conditions (Fig. 6C and D).

**Fig 6.**
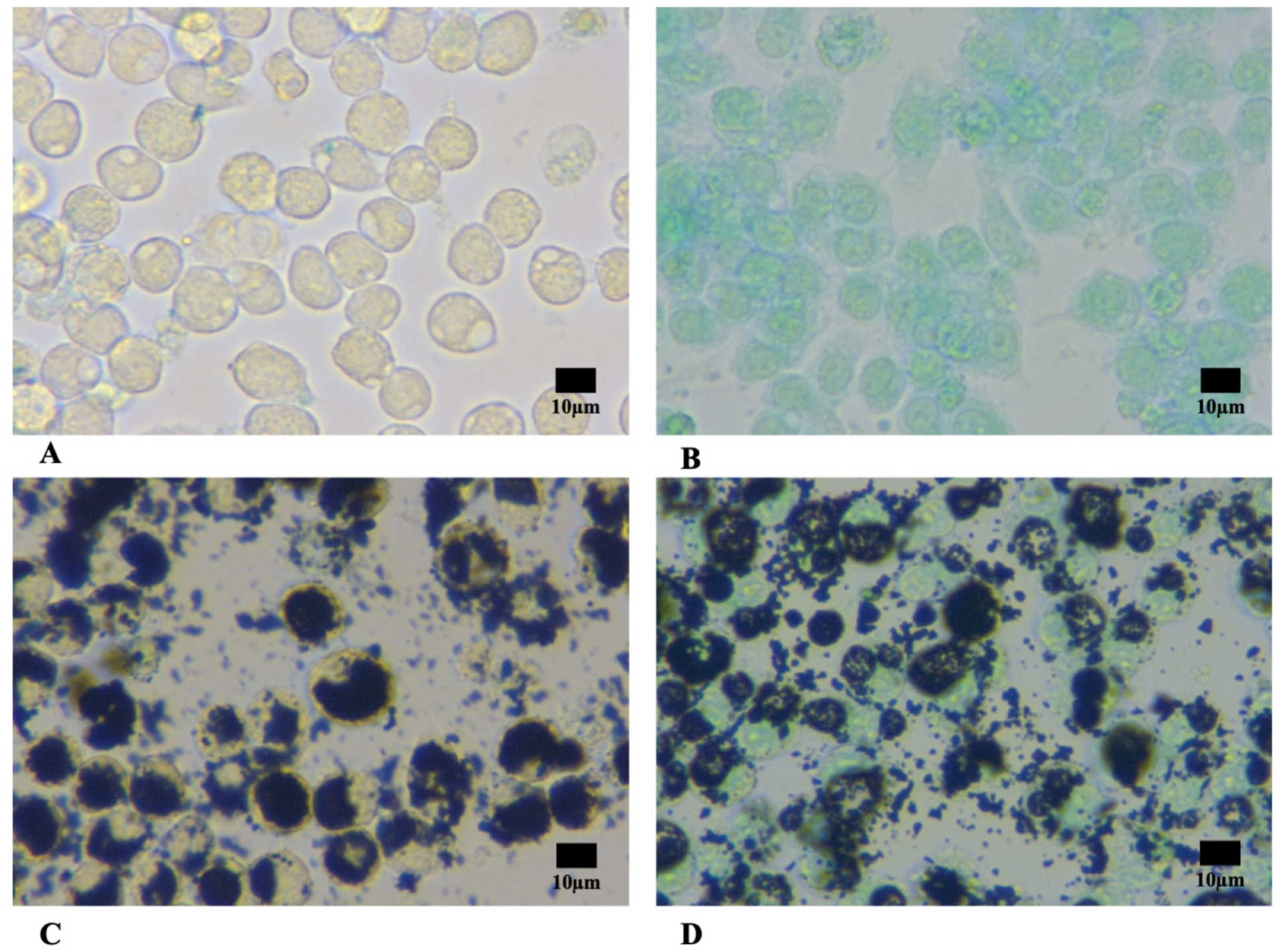
– Viability assessment of J774A.1 cells using VVBlue assay in the presence of colored particle dispersions. Optical microscopy images of live (A and C) or ethanol-killed (B and D) macrophages exposed (C and D) to 1000 µg/ml Pigment black 33. Images were taken with an optical microscope (Motic AE21) at a magnification of 40x. The images were captured with an Axiocam 105 color camera and processed uniformly using ImageJ to standardize contrast and saturation levels.

This robustness of the VVBlue assay allowed us to determine the toxicity curve of the two manganese-based pigments Pbk14 and PBk33. The results, displayed on **Figure 7**, showed that PBk33 was not toxic at concentrations up to 1 mg/ml, while PBk14 became toxic at concentrations as low as 100µg/ml.

**Fig 7:**
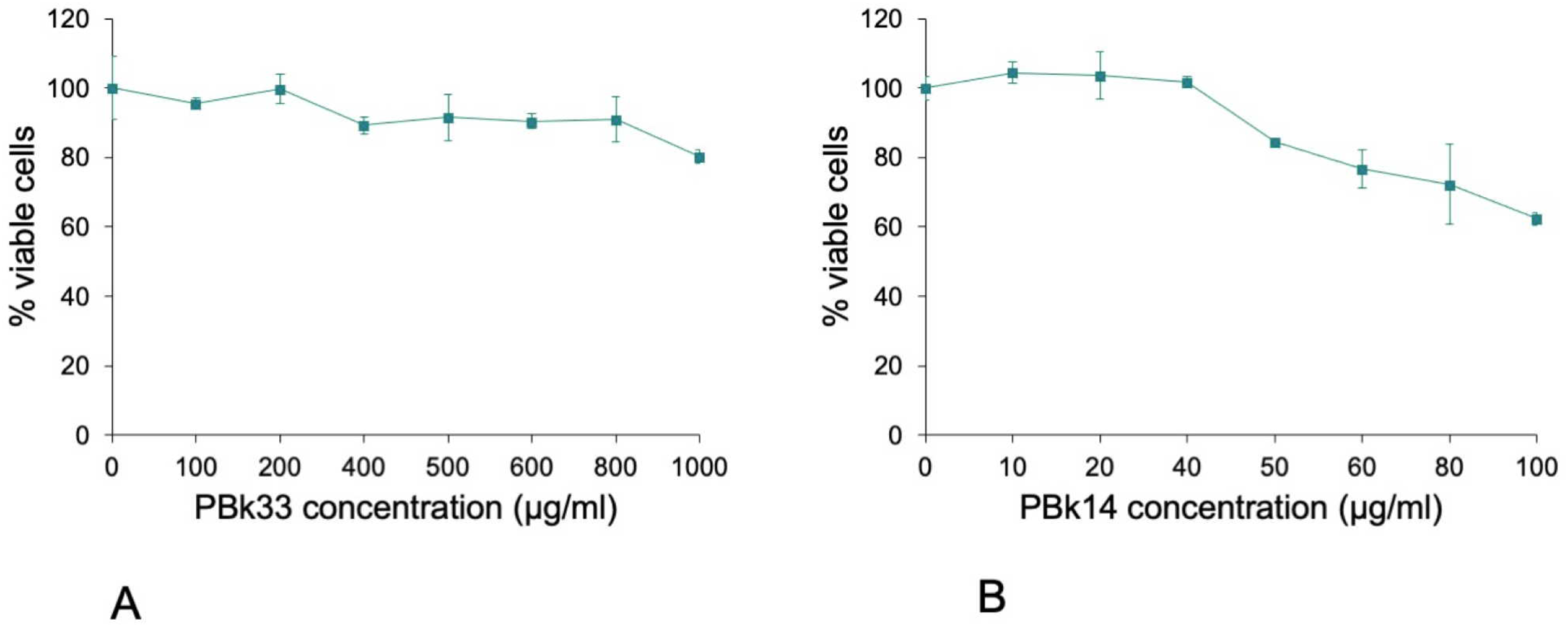
Toxicity curves of two manganese-based pigments. A) Cell viability after 24 hours of exposure to PBk33 (mixed manganese iron oxide). The abscissae are the concentration of pigment added to the cultures in µg/ml B) Cell viability after 24 hours of exposure to PBk14 (manganese dioxide). The abscissae are the concentration of pigment added to the cultures in µg/ml Experiments conducted in triplicates and expressed as mean ± standard deviation

Thus, the VVBlue assay was able to work under conditions where no other dye exclusion method could work, while avoiding the use of indirect assays and keeping cell integrity as the cell viability parameter.

Finally, we wondered whether the VVBlue assay would still work with blue or green pigments, which may interfere with the reading of the alphazurine absorbance. We thus tested the toxicity of the recently discovered Pigment Blue 86 [31], which is authorized for commercialization under a low volume exemption [32]. As the pigment deposited at the bottom of the cell culture wells and formed an opaque blue layer, that stayed at the bottom of the wells, the eluates from the alphazurine and crystal violet staining steps were transferred to a clean reading plate before absorbance reading, as described in the protocol, removing any interference fro the pigments. Control experiments with the pigment alone without cells showed negligible absorbance at 637 nm, comparable to the one of control live cells untreated with the pigment but treated with alphazurine. In these experiments, we compared the viability data obtained via the indirect MTT assay [9] and the ones obtained via the VVBlue assay. The results, displayed on **Figure 8**, showed once again a discrepancy between the results obtained via the MTT metabolic assay and the membrane permeability VVBlue assay.

**Fig 8:**
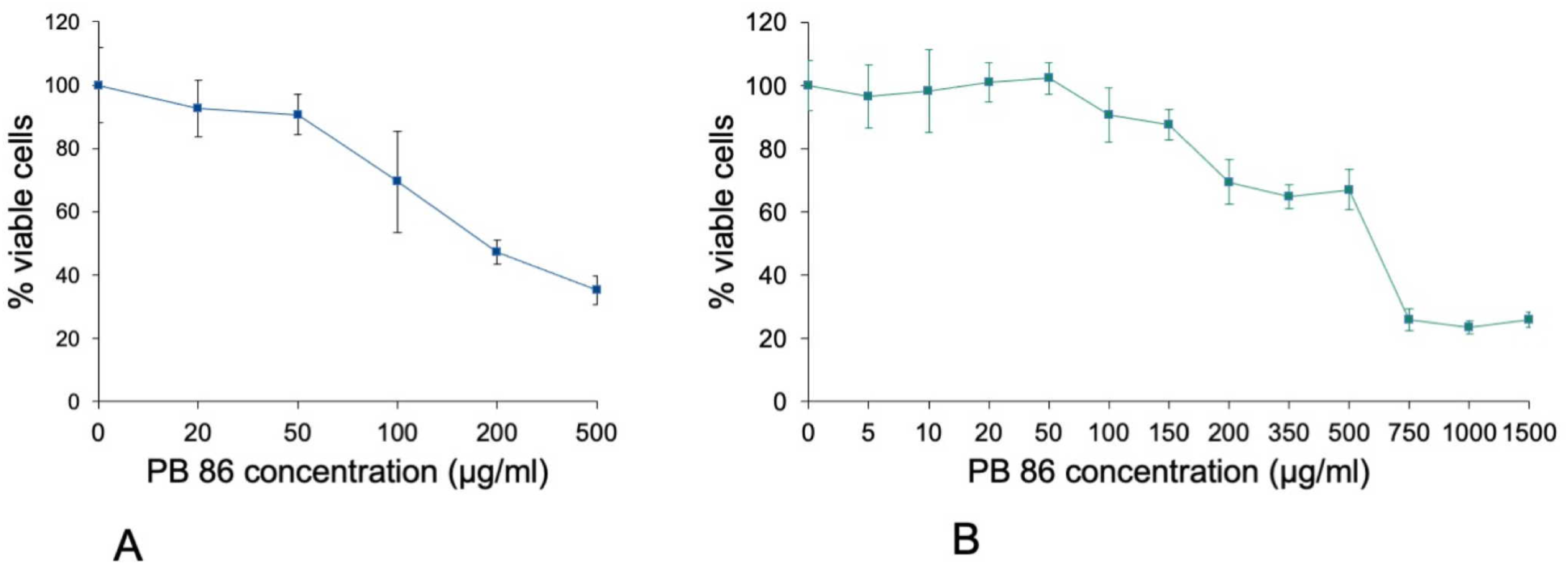
Comparison of the MTT and VVBlue assays on J774A.1 cells exposed to Pigment Blue 86. Cell viability curves after 48 hours of exposure to Pigment blue 86. The abscissae are the concentration of pigment added to the cultures in µg/ml A) data obtained with the MTT assay B) data obtained with the VVBlue assay Experiments conducted in quadruplicates and expressed as mean ± standard deviation

The MTT assay led to a letal dose 20% of 75 µg/ml and a letal dose 50% of 185 µg/ml, while the VVBlue assay led to a letal dose 20% of 150µg/ml and a letal dose 50% of 600 µg/ml. Furthermore, the different shapes of the survival curves were consistent with the mechanisms underlying the two assays. The metabolic assay showed a gradual decrease of the metabolism with increasing pigment concentrations, while the cell permeability assay showed an extended survival followed by a rapid drop in cell viability with increasing pigment concentrations.

## 4. Discussion

In viability assays, it is necessary to interpolate numerical values from the experimental points, as the desired parameters (e.g. LD20, LD50) may not be directly reached from the actual data points. To this purpose, regression analysis and most often linear regression analysis is most commonly used.

In regression analysis, the R² coefficient of determination serves as a statistical measure of how well the regression predictions approximate the actual data points. A close-to-1 R² indicates that the model predicts the biological data accurately, while a slightly lower R², within the range of 0.8<R²<0.91, suggest a close approximation to real data points.

Prior studies on the correlation between cell numbers and absorbance of bound dyes have revealed linear relationships between these two variables [33]. It has been ascertained that crystal violet staining may exhibit a plateau phenomenon at high cells numbers. In this study, distinctions between the two employed dyes have been observed. In contrast to the alphazurine staining, crystal violet displays a curvilinear regression pattern. Upon splitting the crystal violet dot chart into two segments, distinct linear trends emerge, each characterized by different correlation coefficients. The first segment, showing the absorbance of CV over the range from 0 to 10^5^ cells, reveal a correlation coefficient of 0.9981. The second segment, spanning from 2 x10^5^ to 10^6^ cells, reveal a lower correlation coefficient of 0.9694. Despite the slightly diminished R² value in the latter linear phase, it attests to a certain correlation between absorbance and cell numbers within range.

When developing a new assay, it must be standardized against established reference assays to ensure that its reliability and accuracy, and thus its robustness. To this purpose, we evaluated our new method against a validated assay of a similar type (i.e. dye exclusion), namely the propidium iodide (PI) viability assay. The VVBlue assay demonstrated a good correlation with the propidium iodide method, although always indicating higher LD20s. This observation can be attributed to the fact that cell detachment of the cells from the plates was required in the PI method. Indeed this process can induce artificial cell death, especially for very adherent cells such as macrophages that can get damaged upon the harsh conditions required to detach them from the culture plates, and are of course even more fragile when their viability starts to be compromised.

When comparing the results of the VVBlue assay with those obtained with a classical metabolic test (WST1), important differences in the toxicity curves were observed. The same type of discrepancy between metabolic tests and leakage-based ones has been described in the literature [30], with different formats (MTT and LDH leakage in [30] compared to WST1 and Alphazurine A here). Such apparent discrepancies shall not come as a surprise, and are just linked to the indirect nature of the metabolic tests. To give a stylistic example, a halved signal in a metabolic test may indicate either that all the cells are still viable nut with a 50% reduced metabolism, or that half of the cells are dead while the other half exhibit a full metabolic activity. Conversely, if cells become more metabolically active to fight a stress that is not lethal, viabilities above 100% can be obtained, as previously described in the literature [13,14]. Thus, indirect test such as metabolic test or neutral red uptake [19] should be taken with caution in their interpretation.

It can also be noted from Fig. 4 that different cell lines can show different responses to the same toxic, in this case silver ion. Silver ion has been shown to be quite toxic toward J774A.1 cells, where it induces strong oxidative stress and metabolic poisoning [35]. Thus, the rate of poisoning and thus the toxicity may depend on the metabolic status of the cells. Indeed, we observed that the J774A.1 cells consumed their culture medium much faster than the confluent A549 cells that we used, indicating a higher glucose consumption. Macrophages are known to utilize extensively glycolysis, e.g. for phagocytosis [36]. As silver ion is known to inhibit key glycolytic enzymes such as glyceraldehyde phosphate dehydrogenase [37], triose phosphate isomerase [38], phosphoglycerate dehydrogenase [39] or lactate dehydrogenase [40], this may explain the higher sensitivity of macrophages to silver ion.

However, it must be noted, for example on the Pigment Blue 86 case, that the data obtained by the metabolic assay and the cell permeability assay fit well with what is known about the toxic mechanisms shown by metals. Indeed, pigments gradually dissolved when internalized by cells, and release free metal ions, which then bind to some proteins active sites and block their functions [41]. This applies to metabolic enzymes and leads to a decrease in metabolism that is detected by the tetrazolium-based assays (MTT or WST). However, cells are able to survive with a hampered metabolism, up to the point were they cannot compensate any longer and cannot maintain the integrity of their membrane, leading to cell death detection by the permeability assay.

Our experiments also provide the first public data for the toxicological analysis of Pigment Blue 86, a new metal-based pigment [31] that has received limited market authorization [32]. This pigment has limited toxicity, which is worth noting.

Overall, the VVBlue assay represents, to our knowledge, the first exclusion dye assay that is plate-readable. Classical dye exclusion dye methods (e.g. Trypan blue) require either manual reading under a microscope, which is tedious and makes the assay prone to inter-operator variability, or an expensive and dedicated camera type setup. The PI assay requires a flow cytometer, and operator-defined delimitation of the threshold values for live and dead cells, which may be questionable too. At the expense of additional internal controls, namely “live cells” and “dead cells”, the VVBlue assay provides an operator independent reading of the cell viability, ideally using a versatile plate reader instrument. Alternatively, the absorbance of each well can be read individually in a spectrophotometer, although this will severely limit the throughput of the assay.

Furthermore, this assay can operate under conditions in which many of the other cell viability assays become impracticable, as demonstrated here for pigment particles. Of course, further control conditions needed to prevent data misinterpretation caused by dye-particle adsorption issues, but this can be easily implemented, as shown in the tests displayed in Figure 8.

It must be kept in mind that the VVBlue assay reads the number of dead cells in the well. Depending on whether the centrifugation step can keep all the dead cells in the well or not, or if the assay is conducted under conditions where the cells proliferate, the second crystal violet step, which provides the total number of cells still attached to the plate, may be useful or not. In any case, the double reading proposed here covers all situations, and may help distinguishing cytostatic effects from cytotoxic ones.

In conclusion, the VVBlue assay offers the advantages of dye exclusion assays under a plate-readable format. When the adequate controls are added to standardize the assay, it offers a reliable of the cell viability, even under severe optical conditions. However, it also shows some downsides. First of all, because of its mechanism, the alphazurine assay must be coupled with a cell enumeration-based assay, crystal violet assay in our case. This means in turn that the overall assay consists of two successive individual assays, which renders the global assay slow and somewhat cumbersome. Although we have not yet found conditions where the assay becomes ineffective, it can be predicted that toxicants that are able to release a blue or green color in the elution medium will strongly interfere with the alphazurine assay.

Last but not least, as every assay based on numbers and operating in the plate format (such as metabolic assays, neutral red assay etc), and opposite to assays operating on cell suspensions,(such as propidium iodide or trypan blue assays) which operate directly as proportions, the VVBlue assay will be sensitive to cell proliferation, which may increase cell numbers in some conditions and thus bias the toxicity results. Nevertheless, we hope that the VVBlue assay will find utility in determining chemicals toxicities, especially when adverse optical conditions are encountered.

## Data availability

All the data underlying this worked are available in the BioStudies database [42], under the doi: 10.6019/S-BSST1506

## Safety concerns

Acetic acid is an irritant

Ethanol is toxic at high doses

Crystal Violet is a suspected carcinogen and is toxic for the environment

All solutions and wastes used in this protocol should be disposed of respecting applicable local regulations

## Materials

Culture plates shall be with a transparent flat bottom and treated for mammalian cell adherence. We use 12 or 24 well culture plates from Falcon (#353043 and #353047, respectively)

### Chemicals

Non-denatured ethanol should be used. 96% ethanol is acceptable without correction for titer

Acetic acid is 99% pure. We use reference A6283 from Sigma Aldrich

Alphazurine A was reference A2770 from Sigma Aldrich. This product is discontinued. Alternate references are #ICNA0215473025 from MP biomedicals or reference AC189370100 from ThermoFisher.

Crystal Violet is reference C0775 from Sigma Aldrich.

PBS, calcium and magnesium free, is reference D5652 from Sigma Aldrich

## Equipment

A plate reader able to read absorbance is required. The required wavelengths to be read are 590 nm (Crystal Violet) and 637 nm ( Alphazurine)

## Test Setup

The rationale of the alphazurine stain (staining of dead cells in the plate) means that the absorbance reading at 637 nm will provide an index of the number of dead cells. This index must be scaled against wells that will contain only dead cells and thus represent a 100% dead cells level. This means that a plate designed for a complete test shall be arranged as follows

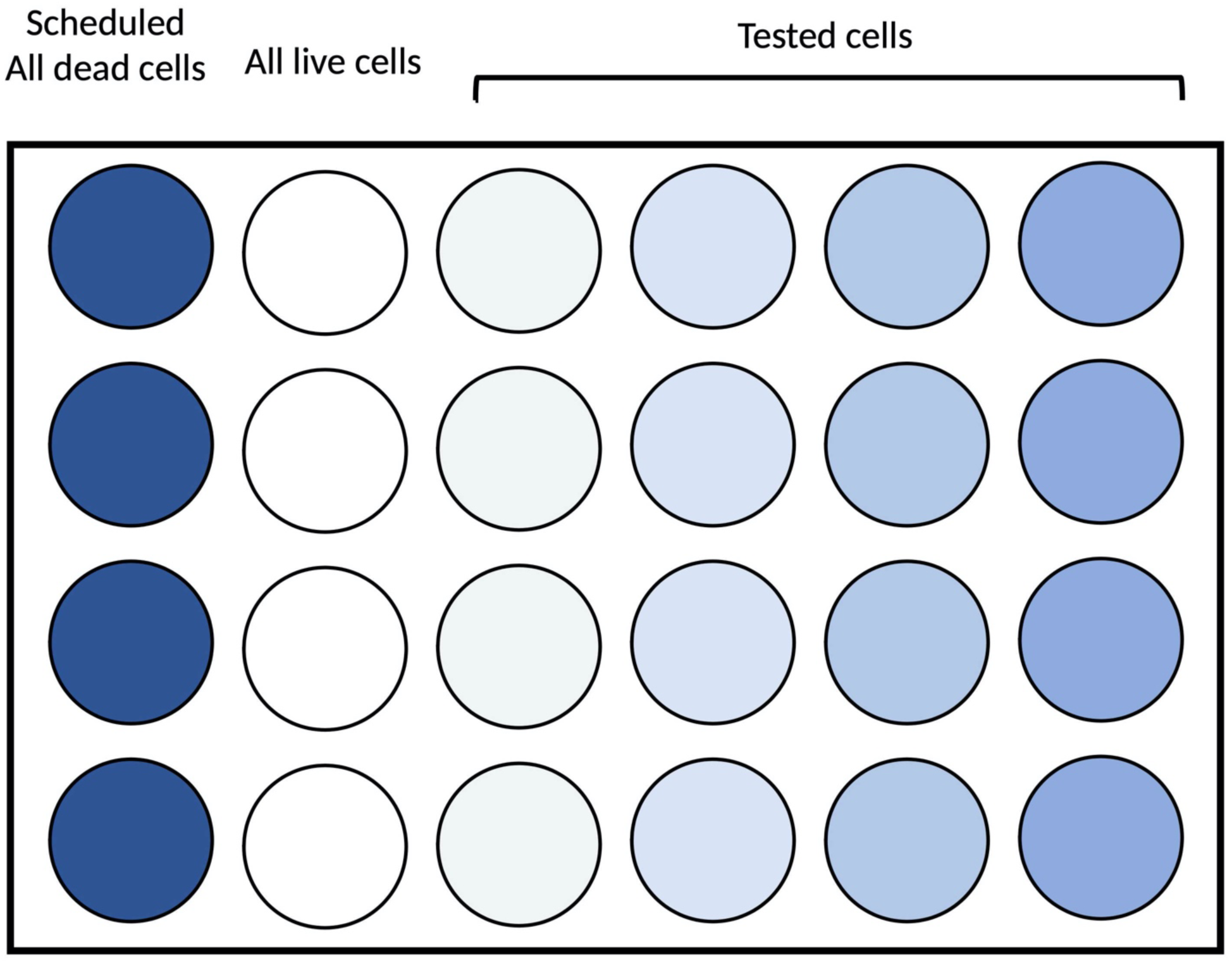
Note: we do not recommend to go below four replicates by condition, which is an absolute minimum.

## Protocol

|  |  |
| --- | --- |
| *1) Plate preparation* |  |
| Centrifuge cell culture plate for 2 min at 1200 rpm | Centrifugation prevents any loss of cells |
| Remove the medium from well scheduled for 100% dead cells control<br>add 300µL of Ethanol (EtOH) 50% for 3-5 min at RT | The addition of ethanol induces cell death and makes their membranes permeable |
| Rinse with DMEM live cell wells | The medium may have been affected by the 48h culture period or the presence of particles, prompting its replacement with a fresh one |
| Remove EtOH 50%<br>Rinse wells once with clean culture medium<br>Add a volume of clean culture medium | The exposure to ethanol dehydrates the cells, necessitating hydration before dyeing them |

| **2) Alphazurine dye staining** |  |
| --- | --- |
| Add AlphazurineA at a final concentration of 0,2 mg/mL in all wells (50 fold direct dilution from a 10 mg/ml in water) | Alphazurine is added to fresh medium to maintain cells in a favourable environment while staining, ensuring that viability remains uncompromised |
| Incubate 40 min at 37°C | Incubating cells under cultured conditions ensures that viability remains uncompromised |
| Centrifuge 3 min at 1200 rpm | Centrifugation prevents any loss of cells |
| Rinse 3x with 1X Phosphate-buffered saline (PBS)<br>Centrifuge for 3 min at 1200 rpm after each rinse | Rinsing cells with PBS facilitates the removal of dye that have not entered the cells<br>Centrifugation prevents any loss of cells |
| Remove the supernatant | The last rinse with PBS has to be remove before adding the elution solution |
| Elute by adding a volume (equivalent to cell culture medium used) of 50% ethanol 1% acetic acid solution | Adding acetic acid to ethanol enhances the dye solubility |
| Incubate 30 min at RT on a shaker<br>Centrifuge for 2 min at 1200 rpm | Centrifugation prevents any loss of cells |
| (Optional) Transfer 200µL into a 96-well plate | To eliminate the potential interference from cell absorbance on the original plate, it is advisable to transfer a portion on supernatant into a new plate |
| Read absorbance at 637 nm |  |

| **3)Crystal violet dye** |  |
| --- | --- |
| Remove the remaining alphazurine eluate<br>Rinse once with elution solution, then once with PBS | The elution solution dehydrates the cells, necessitating hydration before dyeing them with another dye |
| Add one volume (equivalent to cell culture medium used) of CV solution at 4 µg/ml in PBS | As the cells are dead, exposure to water poses no risk to them, so this solution can be prepared using either water or PBS |
| Incubate 30 min at RT | It is not necessary for cells to be under culture conditions for this staining, as all cells are dead |
| Centrifuge 3 min at 1200 rpm | Centrifugation prevents any loss of cells |
| Rinse 3x with PBS<br>Centrifuge for 3 min at 1200 rpm after each rinse | Rinsing cells with PBS facilitates the removal of dye that have not stained the cells<br>Centrifugation prevents any loss of cells |
| Remove the supernatant | The last rinse with PBS has to be remove before adding the elution solution |
| Elute by adding a volume (equivalent to cell culture medium used) of 50% EtOH 1% acetic acid solution | Adding acetic acid to ethanol enhances the dye solubility |
| Incubate 45 min at RT on a shaker<br>Centrifuge for 2 min at 1200 rpm | Centrifugation prevents any loss of cells |
| (Optional) Transfer 200µL into a 96-well plate | To eliminate the potential interference from cell absorbance on the original plate, it is advisable to transfer a portion on supernatant into a new plate |
| Read absorbance at 590 nm |  |

## Readout

The alphazurine stain provides an index of the number of dead cells. Thus, in addition to the desired measurements, the plate(s) must contain wells with all live cells (or the best approximation thereof) and wells with only dead cells. There are therefore three sets of measurements in the plate(s).

A637 _all live cells_

A637 _all dead cells_

A637 _tested cells_

The A _all live cells_ values can be used as a background value, as it integrates the absorbance due to stain adsorbed onto plastic, absorbance of the plate plastic itself and the absorbance due to the few dead cells that may exist in a healthy cell culture.

Then the 100% scale can be calculated with the formula A637 _all dead cells_ - A637 _all live cells_

Then, for each tested condition, the fraction of dead cells in the well can be calculated by the formula Fraction_dead_ = (A637 _tested cells_ - A637 _all live cells_)/ (A637 _all dead cells_ - A 637_all live cells_)

However, these fractions also depend of the total number of cells that are present at the end of the alphazurine reading. Depending on how the tested chemicals act on cells, dead cells may still be adherent to the plate (which is the case for e.g. ethanol) or dead cells can detach or disintegrate and thus be no longer present in the plate when the alphazurine absorbance is read. To compensate for this phenomenon, the crystal violet staining will provide an index of the total number of cells (either initially dead or alive) that are still present at the bottom of the well.

Thus, the fraction of cells present in the well is given by the formula

Fraction _present_ = (A590 _tested cells_ / A590 _all live cells_)

This allows to calculate the fraction of live cells compared to the initial cell input, which is the required value

Fraction _live_= Fraction _present_ – Fraction_dead_

